# Benchmarking of open access publication rates for the pharmaceutical industry and research-intensive academic institutions: development and demonstration of a live, open monitoring tool

**DOI:** 10.1101/2024.09.14.613042

**Authors:** Tomas Rees, Valérie Philippon, Andrew Liew, Slávka Baróniková, William T Gattrell, Jo Gordon, Taija S Koskenkorva, Santosh Mysore, Joana Osório, Tim J Koder

## Abstract

**Background:** In biomedical and health sciences, many articles are published open access (OA). OA publication rates continue to grow, especially for pharmaceutical research. Previous analyses of pharmaceutical OA publication rates were labor-intensive and not easily automated. We aimed to create a free, live, online dashboard comparing OA publication rates between pharmaceutical companies and academic institutions.

**Methods:** Using the Lens – an online platform aggregating full-text content and metadata for over 280 million scholarly works – we built a dashboard showing OA publication rates for medical research articles by authors from the top 40 pharmaceutical companies (pharma) and 40 research-intensive academic institutions (academia). The dashboard details OA rates by model, license and therapy area across three time frames: 12–24 months; 0–12 months; 0–10 years. We downloaded data from the dashboard between 24 July and 4 August 2023 and performed further analysis.

**Results:** Of the articles published 12–24 months before data extraction, 76.6% by pharma and 69.5% by academia were OA. The most common OA models were gold (pharma, 37%; academia, 41%) and hybrid (pharma, 22%; academia, 11%). Oncology had lower rates of OA articles than other therapy areas. In the 10 years before data extraction, growth in the OA publication rate was generally more rapid for pharmaceutical companies than for academic institutions, regardless of field (change in overall % OA 2013–2023: pharma, 3.1; academia, 1.6).

**Conclusions:** This dashboard provides novel and regularly updated evidence on the comparative OA publication practices of pharmaceutical companies and academic institutions. In our snapshot analysis, the OA publication rate was higher for pharmaceutical companies than for academic institutions and continues to increase. Notable differences were observed in approaches, in terms of licenses and OA models, which may reflect different institutional practices and circumstances. We encourage others to use this open resource for new research and to report their results.

## INTRODUCTION

The global pharmaceutical industry is a major funder of healthcare research, with a $238 billion spend on research and development (R&D) in 2021 (Statista 2022). Pharmaceutical companies, which develop and manufacture most medicines, are increasingly interested in having their roles as research funders recognized and in developing trusted reputations similar to those of public and philanthropic funders (Smith 2017).

Open access (OA) refers to the practice of making research outputs, including peer-reviewed articles and data, freely available to everyone. There are several models through which peer-reviewed articles can be published OA, including: bronze OA, which refers to articles that are immediately free to read but do not have a clearly identifiable copyright license; gold OA, in which journals make articles available to read immediately without subscription in exchange for a fee known as an article processing charge (APC); green OA – also called self-archiving – which allows authors to make accepted manuscripts available to read via a personal, funder-controlled or independent repository (immediately or after a defined embargo period); and hybrid OA, which refers to articles that are published in subscription journals but become available to read immediately without subscription through payment of an APC by the authors. In the hybrid OA model, journals collect both subscription fees from individual readers and institutional libraries, and APCs from authors.

Articles published through gold, green or hybrid OA are typically subject to a copyright license (e.g., a Creative Commons [CC] license), which dictates how the published content can be distributed and reused. Some licenses allow anyone to reuse and modify content (e.g., CC BY [Attribution]), while others restrict commercial use (e.g., CC BY-NC [Attribution-NonCommercial]). ‘Commercial use’ is often interpreted to include use by pharmaceutical companies, even if the work will be used for non-promotional purposes. In such cases, commercial organizations must request copyright permission from the journal to reuse materials, which, if granted, can incur a fee.

The OA model and CC license type used to publish research articles vary widely and may be influenced by the country or institution in which the research took place, the funder of the research, the research environment or discipline, researcher perceptions of OA publishing, the journal the article is published in and the authors’ ability to pay APCs (Ayeni & Larivière 2025).

Publishing OA is associated with many benefits for authors, research funders and society. By increasing their OA publication rate, research producers, including the pharmaceutical industry, can build trust in their research by allowing timely scrutiny of their outputs and onward knowledge generation. OA articles can also help make access to scientific literature more equitable for medical professionals, scientists, patients, caregivers and other stakeholders outside the privileged circle of people with access to comprehensive journal subscriptions. In the 2022 *Edelman Trust Barometer Special Report: Trust and Health*, lack of health information was the second most common reason for people to feel unable to improve their health (Edelman 2022). With an increasing volume of poor-quality information online (Borges do Nascimento et al. 2022), OA articles could provide access to credible science to all.

OA articles may also have a greater measurable impact than non-OA articles. Although a large randomized study found no difference in the number of citations between OA and non-OA articles (Davis et al. 2008), other analyses have found that OA articles are more likely to be cited than similar non-OA articles, particularly in medicine (Klebel et al. 2025; Yan & Li 2018). Other studies have found that OA articles in hybrid clinical medical journals were more likely to be cited than paywalled articles in the same journals (Saravudecha et al. 2023), and OA healthcare articles were more likely to be cited in news media than paywalled articles (Schultz 2021).

The proportion of articles published OA is growing. In 2021, 48% of all articles indexed in Web of Science were OA, up from 15% in 2000 (Seo 2023). Between 2009 and 2016, articles funded by large pharmaceutical companies were increasingly being published OA (Yegros & van Leeuwenn 2018). This analysis also found that the gap between the proportion of articles from these companies published OA and the baseline proportion of articles from all types of funders published OA had been closing over time, with the proportion of company articles published OA surpassing the baseline in 2016 (Yegros & van Leeuwenn 2018). Across global regions, OA publication rates in 2015 were highest in sub-Saharan Africa, followed by North America, and Latin America and the Caribbean (Iyandemye & Thomas 2019).

Funder policies can have a key role in influencing OA publication rates, and many funders now have policies that encourage or mandate OA publishing. For example, under the OA mandate of Horizon 2020 (the European Union’s research and innovation funding program from 2014 to 2020), funding beneficiaries have to ensure that peer-reviewed articles arising from the funded work are published OA (European Commission). In 2022, The White House Office of Science and Technology Policy in the USA released renewed policy guidance for government departments toward “ensuring free, immediate, and equitable access to federally funded research” (Nelson 2022; White House Office of Science and Technology Policy 2023). Plan S – a consortium of non-commercial funding agencies, including national funding bodies and independent philanthropic organizations – mandates that the research it funds must be published in OA journals, on OA platforms, or made immediately available through OA repositories without embargo (cOAlition S). Such mandates can instigate great changes in the adoption of OA practices (Huang et al. 2020). Although authors bear the responsibility of complying with OA mandates, for them to do so, journals must offer authors suitable OA publishing options regardless of their funding source. The inclusion of funder policies in directories such as the Registry of Open Access Repository Mandates and Policies (ROARMAP) and Open Policy Finder (formerly Sherpa services) (Jisc) can encourage journals to align their publishing policies with funder requirements. As a result, non-commercial research funders such as Plan S with strong, public OA policies can benefit from more open publication licenses than those without a strong, public commitment to OA (Lancet 2024; NEJM 2024). For academic and non-profit organizations, publishing OA means that the results of publicly funded research are publicly available. Given that the pharmaceutical industry manufactures medicines under licenses granted by governments, it has been argued that the social contract between pharmaceutical companies and general society includes the responsibility for pharmaceutical companies to provide quality information about their products (Edelman 2022). This includes transparently and promptly sharing the results of the extensive R&D they carry out throughout the drug candidate and product lifecycle, and, more specifically, publishing this information OA (Doctorow 2012). The *Good Publication Practice Guidelines for Company-Sponsored Biomedical Research: 2022 Update* (*GPP, 2022*) states that OA options should be prioritized when selecting target journals (DeTora et al. 2022). However, contrary to this goal, some journals do not allow pharmaceutical industry-funded research to be published OA in the same ways in which they allow research funded by non-commercial institutions to be published OA. Further evidence is needed to make the case for the changes to pharmaceutical company publication policies that may influence journal policies (Ellison et al. 2019).

The pharmaceutical industry is increasingly supporting OA publishing, and at least three companies have made public commitments to publish exclusively OA: Takeda (at the time, Shire) in 2018 (Takeda 2018); Ipsen in 2019 (Page et al. 2020); and Galápagos in 2020 (Galapagos 2020). GSK (GSK 2023) and Pfizer (Skobe et al. 2023) have also publicly stated their aims to increase OA publication rates. Furthermore, 18 pharmaceutical companies (including the five mentioned above) have become Members or Supporters of Open Pharma, a multi-sponsor collaboration working to improve the communication of pharmaceutical industry-funded research (Open Pharma). In 2019, Open Pharma released a position statement on OA that outlined the collaboration’s long-term goal to “secure authors publishing company-funded research the same [OA] terms as authors publishing research funded by other sources, so that all research can be made free to read – and reuse – from the date of publication” (Sabir et al. 2019). Other companies in the Open Pharma collaboration have internal policies recommending OA, but details of these policies are not publicly available.

The OA publication rates of different pharmaceutical companies have previously been compared (Koder et al. 2021; Macdonald & Koder 2020), but the method used required substantial manual analysis that was not conducive to automation, and involved the use of proprietary information. Some pharmaceutical companies also analyze their own proprietary records to measure and drive improvement in OA publishing rates (Skobe et al. 2023). However, the methods used differ between companies, and most of the results of these analyses are not available to the public.

Tools such as the live Curtin Open Knowledge Initiative dashboard (Curtin Open Knowledge Initiative; Wilson et al. 2022) and the STM OA dashboard (STM) provide ongoing tracking and comparison of OA publication rates by country. However, to our knowledge, no study has compared the OA publication rates of pharmaceutical companies against those of research-intensive academic institutions.

In this article, we describe the development of a free, live dashboard designed to allow benchmarking and comparison of OA characteristics and trends of medical articles published by authors affiliated with pharmaceutical companies and those published by authors affiliated with research-intensive academic institutions. To illustrate how this dashboard can be used, we then present a snapshot of data from the dashboard as a use case.

Portions of this article were previously published as part of a preprint (Rees, 2024).

## MATERIALS AND METHODS

### The Lens

The Lens (Lens.org 2024) is an online platform that aggregates full-text content and metadata for over 280 million scholarly works from primary data sources, including CrossRef, PubMed and OpenAlex. These primary sources are supplemented with additional data from Unpaywall, OpenCitations, the Directory of Open Access Journals, Open Researcher and Contributor iD, the Research Organization Registry (ROR), PubMed Central, and publishers (e.g., Springer Nature) to create ‘MetaRecords’ (Jefferson et al. 2019). Each MetaRecord can be searched by its linked metadata, allowing filtering by categories, including author name, field of study, Medical Subject Headings, institution and funding organization.

The Lens data are updated every 2 weeks, allowing for real-time analysis of trends in scholarly publishing.

### Development of the live dashboard

With support from the Lens, we produced a Lens Report that presents real-time, automatically updated data and analytics related to the OA rates, types and licenses of medical articles with authors affiliated to pharmaceutical companies and research-producing academic institutions (**Fig. 1**). This Lens Report, now called the *Open Pharma open access dashboard*, can be freely accessed here: https://www.lens.org/lens/report/view/Live-open-access-analysis/14572/page/14573. Below we describe the search filters used to produce the dashboard in the Lens.

**Figure 1.**
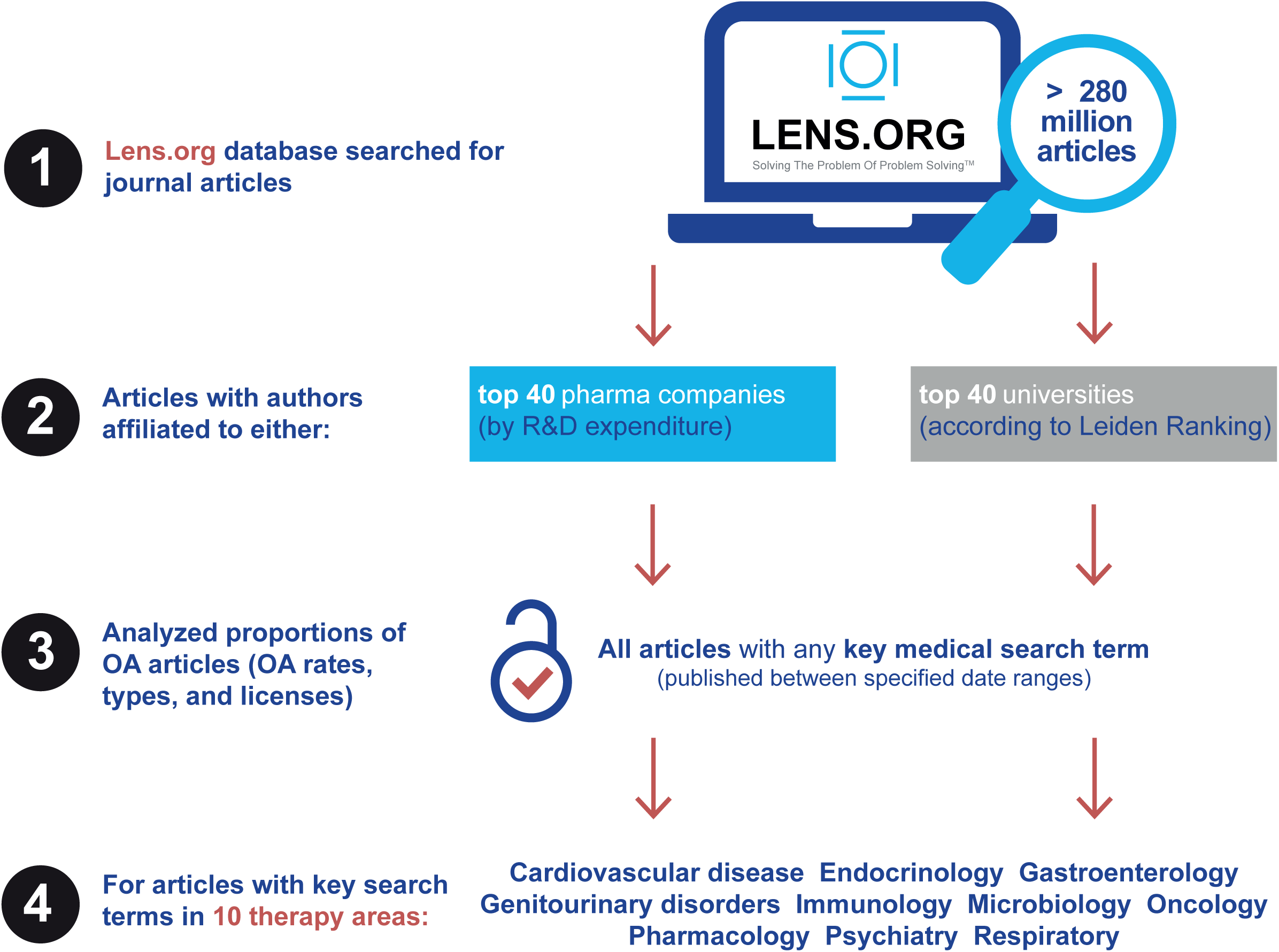
Development of the Open Pharma open access dashboard using the Lens platform.

#### Selection of pharmaceutical companies and academic institutions

First, we filtered the Lens database for journal articles by authors affiliated with the top 40 global pharmaceutical companies and a comparator group of 40 academic institutions.

To identify the top 40 global pharmaceutical companies, we used 2022 global R&D spending as reported by the Statistica *Leading 50 global pharmaceutical companies by prescription sales and R&D spending in 2022* report (**Supplemental Table S1**) (Statista 2023). The lowest-ranking company, Grifols, was substituted with Galápagos, a company with similar R&D spend (Galapagos 2021) but that is also an Open Pharma Member and has a greater publication output. At the time of this analysis (Q3 2023), Open Pharma had 15 Members and Supporters. Of these, 14 were pharmaceutical companies, and all were included in the analysis (**Supplemental Table S1**). At the time of writing (Q4 2024) Open Pharma has 18 Members and Supporters, 17 of which are pharmaceutical companies. All except one of these companies is included in this analysis (**Supplemental Table S1**).

A global comparator group of 40 top academic institutions was identified by analysis of author affiliations in the Centre for Science and Technology Studies Leiden Ranking 2022 (**Supplemental Table S2**) (Leiden University Centre for Science and Technology Studies 2022). The Leiden Ranking was selected over other university ranking scales because it compares universities purely by their publication output and performance. We analyzed academic institutions with articles in the ‘Biomedical and health sciences’ field published in 2017–2020. Countries of affiliation were allocated to one of seven regions (Africa, Asia, Europe, Middle East, North America, Oceania and South America), and the institutions with the highest proportion of articles from each region were selected to give a total of 40 academic institutions that reflected the regional makeup of the full sample. The regional composition of these academic institutions is shown in **Supplemental Table S3**. A fully equivalent regional analysis for pharmaceutical companies is not possible because much of their research is not conducted in the country in which the company is headquartered.

Where applicable, we used the ‘primary’ name of each company or academic institution and the common ‘other name’ from ROR.org as well as the ROR IDs lineage (‘author.affiliation.ror_id_lineage’) and child organization names for companies.

Author affiliations were used to identify articles produced by pharmaceutical companies and academic institutions. This approach was adopted because funding sources are not consistently disclosed in article funding statements, making it challenging to identify all the organizations that financially supported the research presented.

#### Definition of medical articles

The Lens includes ‘field of study’ tagging provided by the Open Alex database. Open Alex uses artificial intelligence parsing of all accessible text to assign each of its > 280 million scholarly MetaRecords a ‘field of study’ tag. There are many thousands of such tags, with the most common being umbrella terms such as ‘Medicine’, ‘Ecology’, ‘Library science’, ‘Marketing’, ‘Pure mathematics’ etc. We excluded all articles that did not have a ‘field of study’ tag that was either ‘Medicine’, ‘Internal medicine’ or ‘Pharmacology’. These articles are defined hereafter as ‘medical articles’. This process resulted in the elimination of academic articles from fields that were not comparable to pharmaceutical company research.

#### Time frame

The live dashboard presents data across three time frames: the period from 12 to 24 months before the date on which the dashboard is accessed (referred to as current date; primary analysis); the 12-month period prior to the current date (secondary analysis); and the 10-year period before the current date.

Some OA articles are embargoed for up to 12 months (Khoo & Lay 2018). The primary analysis (articles published 12 to 24 months before data extraction) allows the comparison of all OA articles, including both immediate and embargoed OA articles.

The 10-year time frame was chosen to allow longitudinal comparison of OA trends over time.

#### Definition of therapy areas

We used the ‘field of study’ tags again to define therapy areas, this time looking at the less common tags. The top 1000 fields of study (by publication count) were identified from the medical-related pharmaceutical company articles 10-year subset. Fields of study related to disease or pharmacology terms were identified and then assigned to therapy areas. For example, articles tagged with “Diabetes mellitus”, “Endocrinology”, “Insulin” or “Type 1 diabetes” etc. were included in the therapy area “Endocrine and metabolic disorders”. Sixteen therapy areas were identified, of which the largest 10 were included as categories in this analysis (**Supplemental Table S4**). The top 5 journals for numbers of articles in each of the 10 therapy areas were identified for pharmaceutical companies and academic institutions.

### Snapshot assessment of OA

After completion of the dashboard build using the parameters outlined above, we extracted a data snapshot. Publication count data were collected between 24 July 2023 and 4 August 2023 via the Lens application programming interface (API) (The Lens) using the ‘httr’ and ‘rjson’ R packages. Sen’s slope was calculated using the R ‘trend’ package to estimate the trends in overall and therapy area OA percentages between 2013 and 2023 (R Documentation ; Sen 1968). OA percentages were based on the total publication count, while OA model percentages were based on the total OA publication count. All analyses were conducted in R. The R code for the Lens API queries and slope calculations is available on GitHub: https://github.com/OxPG-Informatics-and-Data-Science-Team/open-pharma-open-access-analysis.

We used OA status, model (referred to as ‘color’ in the Lens documentation) and license types as reported by the Lens (The Lens). Each article is listed with a single OA status (yes or no) and model (bronze, gold, green, hybrid or unknown) but can be listed with more than one license type if the license for a particular article is reported differently in different sources. To simplify the analysis of licenses, license types were classified as ‘unrestrictive’ (Public Domain [CC 0] or CC BY), ‘restrictive’ (CC BY-NC, CC BY-NC-NoDerivatives [ND], CC BY-NC-ShareAlike [SA], or CC BY-SA), ‘publisher-specific’ (e.g., Elsevier, American Chemical Society) or ‘unknown’ (implied OA or unknown) (**Supplemental Table S5**). For articles with multiple recorded license types, we chose the most restrictive applicable category.

OA rates are likely to be influenced by social and economic factors that vary across the globe. Pharmaceutical industry studies tend to be highly international, involving investigators from many parts of the world, which is why we chose a regionally representative sample of universities as the comparator set. However, as 28 of the 40 pharmaceutical companies included have headquarters in North America or Europe, representing 90% of total industry articles, a sensitivity analysis was conducted with a restricted comparator set created by excluding academic institutions outside of North America and Europe. Data for the primary analysis cut (24 July 2021–24 July 2022) (***North America/Europe sensitivity analysis***) were collected from the dashboard on 20 November 2024.

### Patient and public involvement

Patients were not involved in the conduct or dissemination of this research.

## RESULTS

### Live dashboard

We have created a live, free, dashboard that provides a visual presentation of OA publication rates for the top 40 global pharmaceutical companies and a comparator group of 40 global academic institutions in a continuously updated format. The dashboard allows users to reproduce most of the underlying searches that were developed. The dashboard can be found at the following URL: https://www.lens.org/lens/report/view/Open-access-dashboard/14572/page/14573.

### Data snapshot

Below we present a snapshot of data extracted from the dashboard between 24 July and 4 August 2023. These data compare the OA rates, models and licenses of medical articles published by authors affiliated with the top 40 global pharmaceutical companies and a comparator group of 40 academic institutions.

#### Sample

For the 12-to-24-month time frame, there were approximately 20 times more total articles from academic institutions (351 514) than from pharmaceutical companies (18 461) (**Fig. 2**). When filtered for ‘Medicine’, ‘Internal medicine’ or ‘Pharmacology’ fields of study, proportionally more pharmaceutical company (71.0%) than academic institution articles (52.5%) remained in the data sets for analysis.

**Figure 2.**
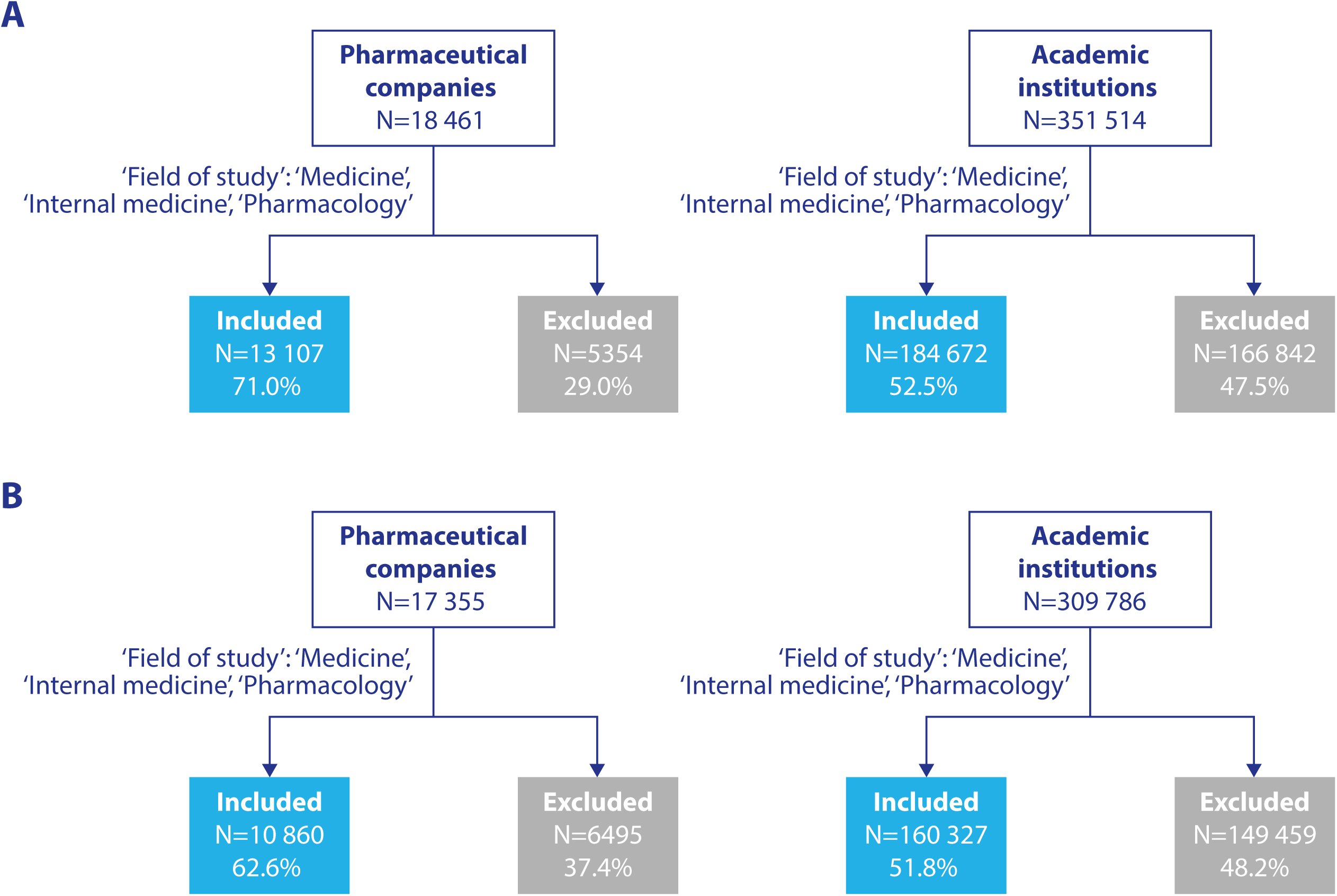
Number of pharmaceutical company and academic institution publications included/excluded in (A) the primary analysis (publications 12–24 months old) and (B) the secondary analysis (publications ≤12 months old).

#### Primary and secondary analyses

Pharmaceutical companies showed a slightly higher proportion of OA articles than academic institutions across both the primary (76.6% vs 69.5%) and secondary (74.4% vs 65.6%) analyses (**Table 1**). A high proportion of articles from both pharmaceutical companies and academic institutions used the gold OA model, but a higher proportion of pharmaceutical company articles than academic institution articles used the hybrid OA model across both analysis periods (**Fig. 3**). In the primary analysis, unrestricted licenses were the most frequently used license category for academic institution articles, while restricted licenses were the most frequently used category for pharmaceutical company articles. In both analysis periods, a greater proportion of academic institution articles had unrestricted licenses than pharmaceutical company articles (**Table 1**).

**Figure 3.**
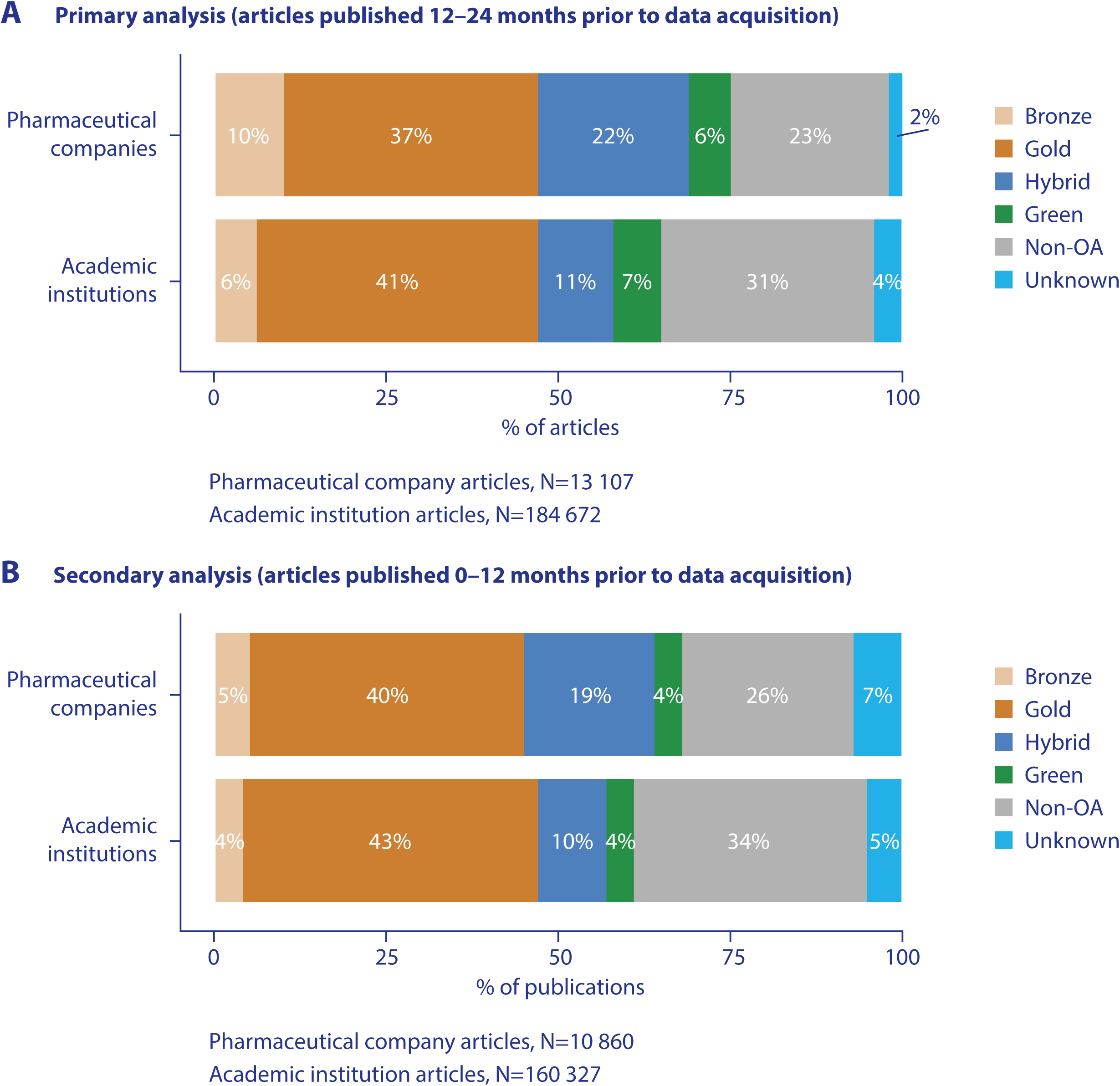
Proportions of pharmaceutical company and academic institution medical publications using different OA models in (A) the primary analysis (publications 12–24 months old) and (B) the secondary analysis (publications ≤12 months old). Percentages may not sum to 100 due to rounding. OA, open access.

**Table 1.** Overall OA/non-OA status and license categories used in pharmaceutical company and academic institution medical articles.

| OA status and license category* | Pharmaceutical company | Academic institution |
| --- | --- | --- |
| **Primary analysis (articles 12–24 months old†)** |  |  |
| Total, n (%) | 13 107 (100.0) | 184 672 (100.0) |
| Non-OA, n (%) | 3066 (23.4) | 56 355 (30.5) |
| OA (any license), n (%) | 10 041 (76.6) | 128 317 (69.5) |
| OA, unrestricted license, n (%) | 2685 (20.5) | 54 147 (29.3) |
| OA, restricted license, n (%) | 3757 (28.7) | 29 929 (16.2) |
| OA, publisher license, n (%) | 2 (0.0) | 122 (0.0) |
| OA, unknown license, n (%) | 3598 (27.5) | 44 189 (23.9) |
| **Secondary analysis (articles ≤ 12 months old‡)** |  |  |
| Total, n (%) | 10 860 (100) | 160 327 (100) |
| Non-OA, n (%) | 2811 (25.9) | 55 232 (34.4) |
| OA (any license), n (%) | 8049 (74.4) | 105 095 (65.6) |
| OA, unrestricted license, n (%) | 2052 (18.9) | 44 160 (27.5) |
| OA, restricted license, n (%) | 2113 (19.5) | 20 514 (12.8) |
| OA, publisher license, n (%) | 2 (0.0) | 89 (0.0) |
| OA, unknown license, n (%) | 3883 (35.8) | 40 397 (25.2) |

#### Trends in OA articles

Over the period 24 July 2013 to 24 July 2023, 63.8% (n = 106 329) of pharmaceutical company medical articles and 62.8% (n = 1 335 346) of academic institution medical articles were published OA. Over the same period, academic institutions saw greater growth than pharmaceutical companies in the overall number of medical articles published (**Fig. 4**). For academic institutions, the proportion of OA articles gently increased over this period from 53.0% in 2013 to 68.4% in 2022 (**Fig. 5**). For pharmaceutical companies, the proportion of OA articles per year also increased over this period from 47.5% in 2013 to 73.6% 2022. Interestingly, the proportion of OA articles published by both academic institutions and pharmaceutical companies appeared to stabilize between 2021 and 2023 (**Fig. 5**). However, the numbers of articles for 2023 were lower than for previous years because data were extracted in July of that year and hence do not represent a full year. Therefore, the proportions of 2023 articles that were OA were likely underestimates because some articles were published under a 1-year embargo after which they will count as OA.

**Figure 4.**
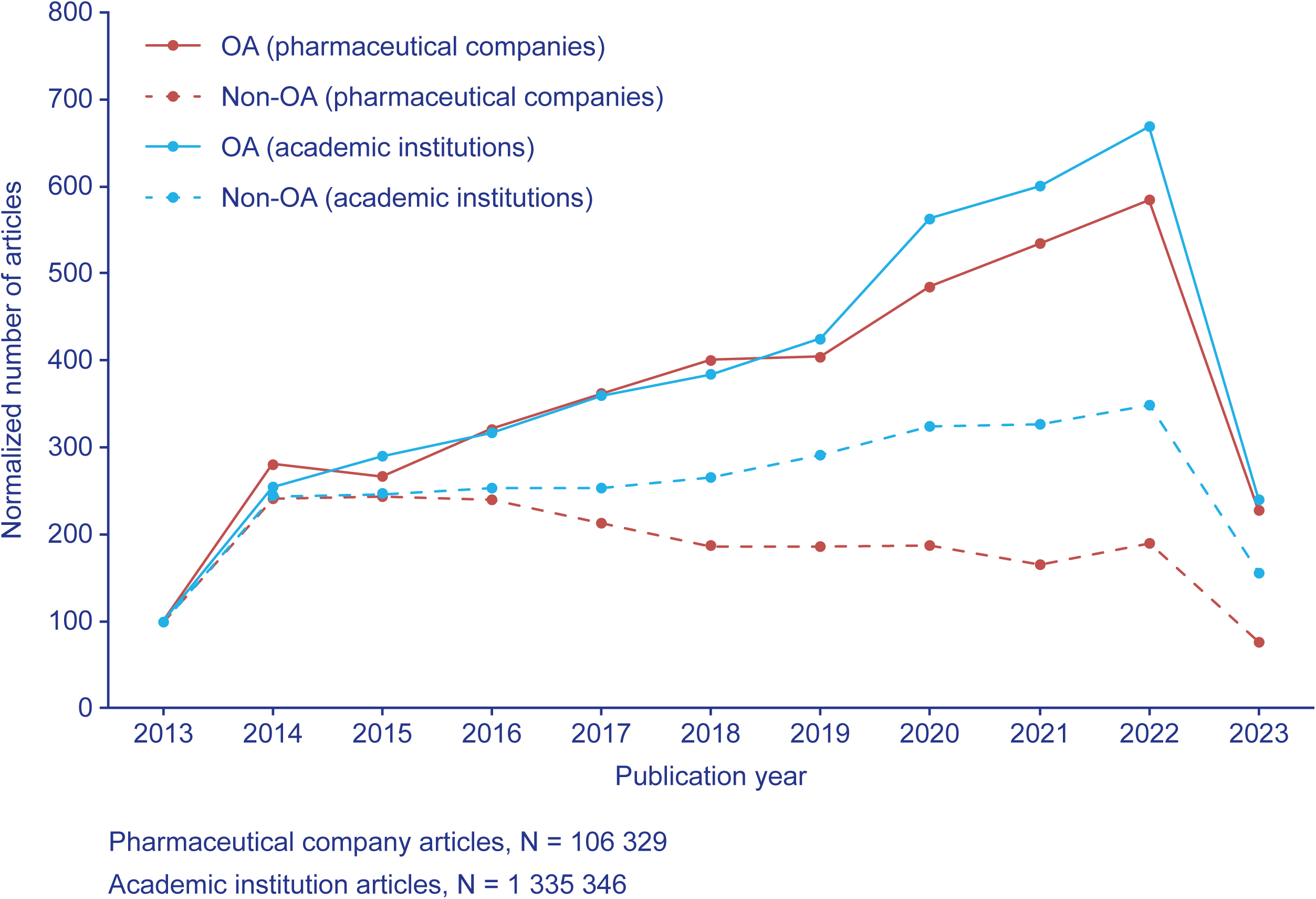
Normalised number of OA/non-OA pharmaceutical company and academic institution medical publications over 10 years (2013–2023). Numbers of publications were normalised to the number of non-OA publications for pharmaceutical companies and academic institutions in 2013. OA, open access.

**Figure 5.**
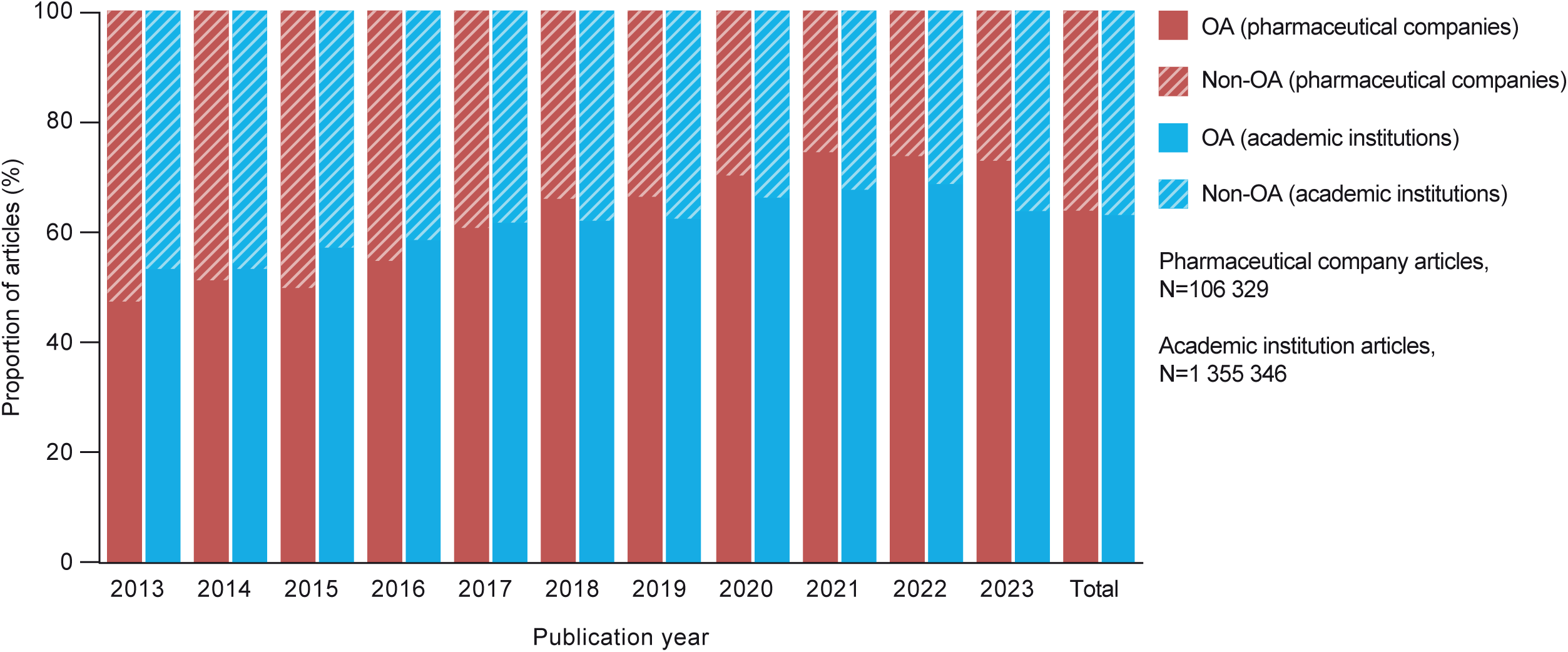
Proportion of OA and non-OA articles in each publication year of pharmaceutical companies and academic institutions.

Regarding OA models, the greatest growth has been in gold OA for both pharmaceutical companies and academic institutions, with gold OA increasing approximately by 1.4% per year and 2.3% per year for pharmaceutical company and academic institution articles, respectively (**Fig. 6** and **Supplemental Table S6**). Pharmaceutical companies also saw an increase in hybrid OA articles of 1.1% per year.

**Figure 6.**
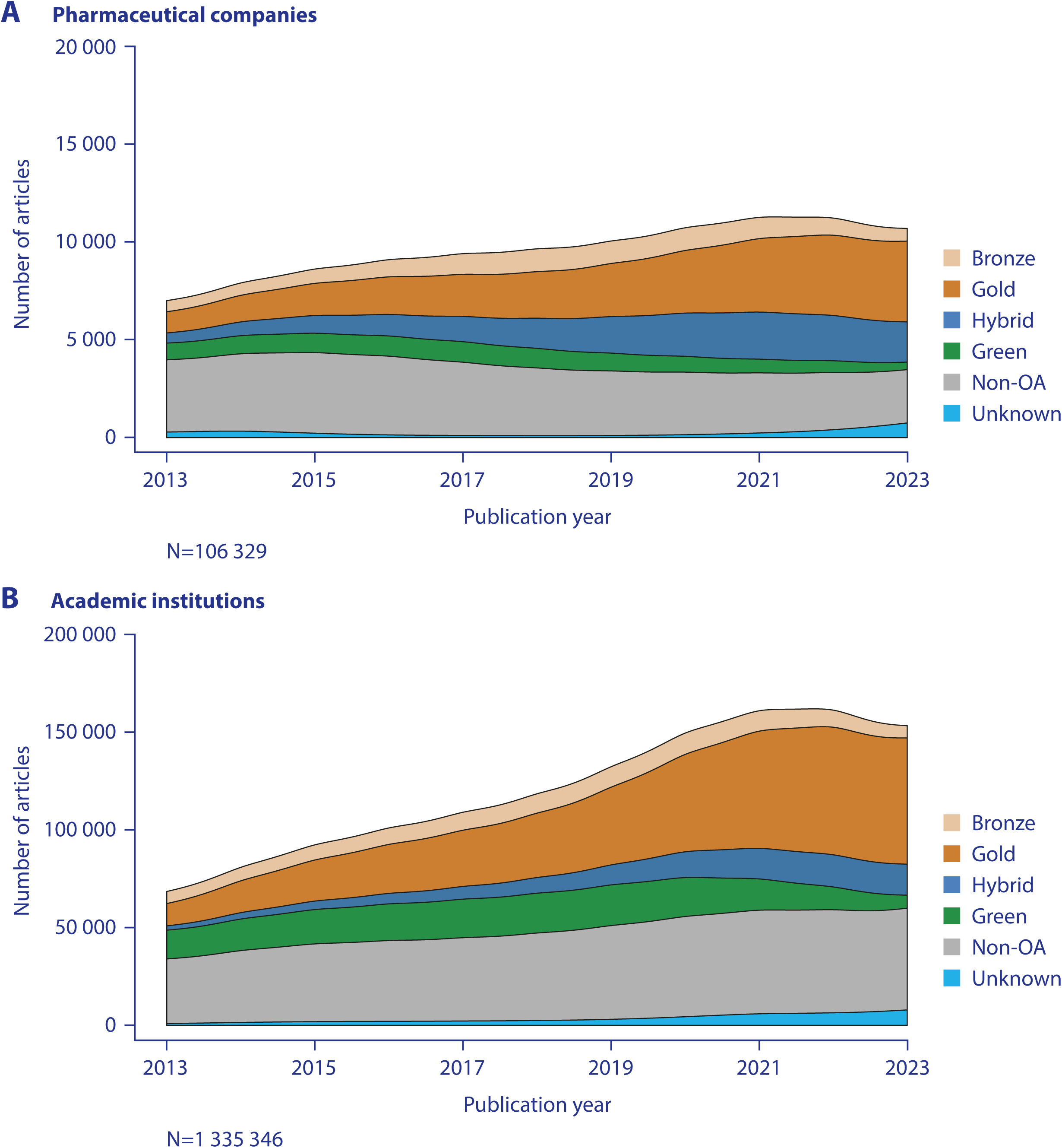
Number of medical publications by OA model over 10 years (2013–2023) from (A) pharmaceutical companies and (B) academic institutions. Data were smoothed in R. OA, open access.

The growth in the proportion of articles that were OA was broadly similar across therapy areas (**Fig. 7, Supplemental Table S6** and **Supplemental Table S7**). Pharmaceutical companies had increases in the proportion of OA articles in all therapy areas. Additionally, growth in the OA publication rate was greater for pharmaceutical companies than for academic institutions in all fields, especially gastroenterology, genitourinary disorders, pharmacology and psychiatry. Oncology, a key research area for pharmaceutical companies, had lower OA publication rates than most other therapy areas, but rates grew year on year.

**Figure 7.**
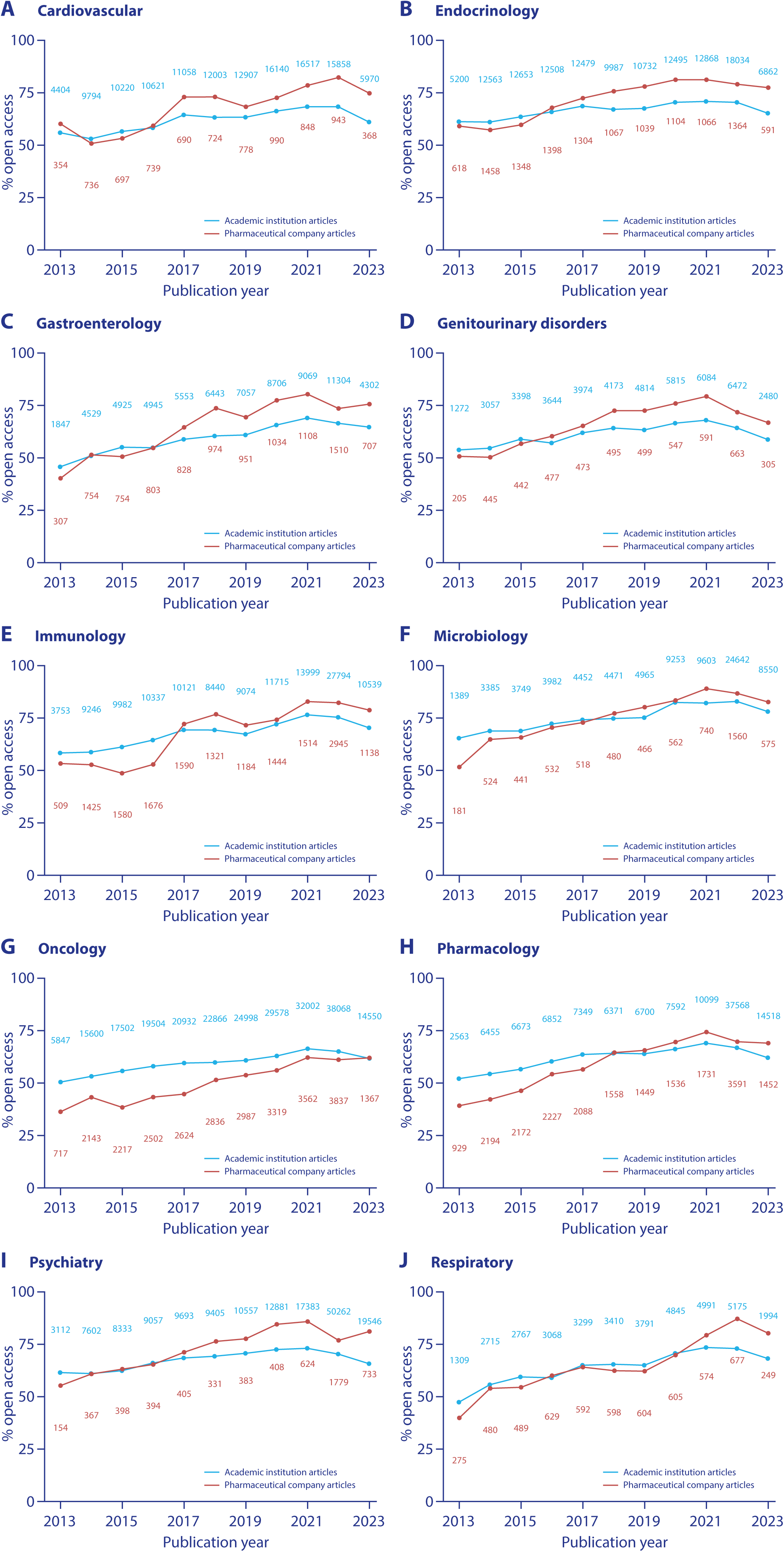
Proportion of pharmaceutical company and academic institution medical publications that are OA over 10 years (2013–2023) by the following therapy areas: (A) cardiovascular, (B) endocrinology, (C) gastroenterology, (D) genitourinary disorders, (E) immunology, (F) microbiology, (G) oncology, (H) pharmacology, (I) psychiatry and (J) respiratory. The number of publications by pharmaceutical companies (red text) and academic institutions (blue text) are shown for each year. OA, open access.

There is substantial divergence between pharmaceutical companies and academic institutions in journal choice. Notably, journals from the natively gold OA publisher Frontiers are popular choices for academic institution authors in several therapy areas, but they do not appear on the list of the top 5 most popular journals for pharmaceutical companies for any therapy area (**Supplemental Table S8**).

#### North America/Europe sensitivity analysis

To account for regional differences OA publishing, we conducted an additional sensitivity analysis comparing the OA rates of pharmaceutical companies with academic institutions in North America and Europe. The data used for this sensitivity analysis, although relating to the primary analysis period (24 July 2021–24 July 2022) were downloaded from the dashboard approximately 16 months later than the primary analysis cut.

Restricting the comparator set to articles from academic institutions in North America and Europe resulted in the exclusion of 38% of academic articles.

Overall, our findings regarding OA models (**Supplemental Table S9)** and license types (**Supplemental Table S10**) remain robust, although the gap in the headline OA rate narrows. A key contributor to the higher overall OA rates for academic institutions in this sensitivity analysis is a relative increase in green OA articles (from 7% to 9% for academic institutions versus a constant 6% for pharmaceutical companies). This may reflect regional differences in self-archiving of articles but may also indicate that articles have been added to green OA archives since our original data extraction.

## DISCUSSION

The development of our OA dashboard represents a substantial step forward in enabling the benchmarking and monitoring of OA publication rates in the pharmaceutical industry and academia. By leveraging data from the Lens, we have created a freely accessible, continuously updated tool that allows users to explore OA trends, licensing models, and journal preferences across multiple time frames. This resource eliminates the need for labor-intensive manual analyses and provides a standardized approach to evaluating OA practices across sectors. The dashboard not only facilitates transparency and accountability in scientific publishing but also serves as a practical resource for researchers, publishers, and policymakers seeking to understand and influence OA adoption.

In our initial snapshot analysis using this dashboard, we show that the OA publication rates of pharmaceutical companies are generally similar to or higher than those of academic institutions. We also show that the total number of articles has grown year on year for both pharmaceutical companies and academic institutions, especially that of OA articles, and that the number of non-OA articles has decreased year by year for pharmaceutical companies but not for academic institutions. Our findings reflect those of Piwowar and Priem (Piwowar et al. 2018), who found that the proportion of journal articles published OA each year increased steadily between the 1990s and 2017. More recently, this trend has been confirmed by Seo (Seo 2023).

The median proportion of academic institution articles (all subjects) published OA during the period 2014–2017 has previously been reported as 43% (Robinson-Garcia et al. 2020), which is lower than the proportion we report for academic institution medical articles (62.8% OA over the period 2013–2023). *Yegros & van Leeuwen (2018)* also found that just over 40% of articles from 2016 with authors affiliated with 23 large pharmaceutical companies were OA (Yegros & van Leeuwenn 2018), whereas our result for 2016 is higher at 54.7%. Additionally, *Yegros & van Leeuwen (2018)* found that articles from pharmaceutical companies were more likely to be published as green OA and less likely to be published as gold OA than articles from all sources (Yegros & van Leeuwenn 2018). This is different to our findings, potentially because we also include bronze and hybrid models in our analysis and potentially also due to the different data sources available in the Lens compared with those used by *Yegros & van Leeuwen (2018)*. However, in agreement with our results, *Yegros & van Leeuwen (2018)* found that the proportion of articles published as green OA decreased between 2009 and 2016, while the proportion published as gold OA increased (Yegros & van Leeuwenn 2018).

### Why are OA rates growing relatively faster for the pharmaceutical industry?

Academic institutions were pioneers in the OA movement. Several large funding bodies have supported and advocated OA for some time, including organizations such as Wellcome (Wellcome), the Howard Hughes Medical Institute (Institute 2024) and the Max Planck Society (Societes 2024), which collectively established the not-for-profit journal *eLife*, further enabling OA publishing. However, in contrast to these large funding bodies, there is likely to be a large amount of academic research funded by smaller organizations for which additional funding for APCs is not readily available. This also contrasts with research funded by pharmaceutical companies. Articles arising from pharmaceutical company research are typically coordinated by specialized publication professionals, a group who are becoming increasingly aware of the benefits of OA and the options for OA articles, and who are increasingly setting and strengthening company OA policies (The International Society for Medical Publication Professionals 2019). The inclusion in GPP 2022 of guidance about prioritizing OA options when selecting target journals is likely to also influence the choices of pharmaceutical company authors and publication professionals (DeTora et al. 2022). Public scrutiny may encourage further adoption of OA practices, and our analysis suggests that pharmaceutical companies are keen to increase their OA publication rates, especially considering the prospect of having the freedom to reuse material from published articles about the studies they fund without needing to seek copyright permission from journals (although this is only possible with permissive copyright licenses such as CC BY). Furthermore, the organizational structure of pharmaceutical companies could mean that OA policies are easier to introduce and implement than similar policy changes would be at academic institutions.

Many academic authors still aim to have their research published in journals with high impact factors despite the movement to reduce the reliance on journal prestige for research assessment (DORA). This is likely to remain the case while academic career progression is based on an individual’s publication record (Ayeni & Larivière 2025). Many of these prestigious journals are subscription journals with limited or no OA options, and academic grants may not cover the higher APCs that are often demanded from more prestigious journals to publish OA (Budzinski et al. 2020).

In the present analysis, we find that OA trends by disease area are similar between pharmaceutical companies and academic institutions, including notably fewer OA articles in oncology than in other therapy areas. This may be because oncology articles reporting key efficacy outcomes from clinical trials are often likely to be targeted to prestigious journals that have more restrictions on OA than other journals. Therapy area may therefore affect the feasibility of implementing OA mandates within pharmaceutical companies, which could be perceived as restricting the authors’ choice of target journals.

Proportionally more pharmaceutical company than academic institution articles are published in hybrid journals, and proportionally fewer pharmaceutical company than academic institution articles are published as gold OA, that is, in fully OA journals. We found that, in some individual journals, the OA publication rates for pharmaceutical companies are lower than those for academic institutions, and we also found that the most permissive copyright licenses are seen less often in pharmaceutical company articles. The lower rates of unrestricted licenses across pharmaceutical company articles are unlikely to be a result of author or company choice, but reflect journals not offering those options for pharmaceutical company research articles (Ellison et al. 2019). These trends can change if journals and publishers choose to allow pharmaceutical companies to publish their articles OA, particularly under the fully open CC BY license allowed to some institutional funders, such as the members of Plan S. Such changes in journal policy are not unprecedented; for example, after engagement with the pharmaceutical industry, the American Society of Clinical Oncology decided in 2018 to allow industry-sponsored research to be published in the *Journal of Clinical Oncology* under a CC BY-NC-ND license after previously not allowing industry research to be published in the journal under any OA license (Ellison 2018). In turn, several journals base their OA offer on the public availability of an organization’s statement of publication policy that includes their policy on OA, and few pharmaceutical companies have made their OA commitments public. Pharmaceutical companies may be in a stronger position than academic institutions to cover the cost of APCs more routinely; however, the increased adoption of read-and-publish agreements may make hybrid OA more attractive for academic institutions in the future (Wenaas 2022).

### Limitations of this analysis

As expected with large-scale bibliometrics sampling, there are many points at which error and/or information loss enter this analysis, including automated affiliations and gray areas on categorizing topics and article types. Inclusion of abstracts wrongly labeled as articles, for example, may underestimate the true volume of OA articles. One journal that was found to be a popular submission target for academic institution articles at the time of the present analysis, *SSRN Electronic Journal*, is in fact a preprint repository, not a peer-reviewed journal. Preprints on *SSRN Electronic Journal* are not persistent and are deleted from the repository following peer-reviewed publication in an Elsevier journal. Our previous OA benchmarking analysis used a more accurate method for article type screening, but it was less conducive to automation than the current method and therefore not suitable for the development of a live dashboard (Koder et al. 2021; Macdonald & Koder 2020). The proportion of articles in *SSRN Electronic Journal* was low, and so this is unlikely to have affected our overall conclusions.

Our set of pharmaceutical company articles essentially includes all articles with a pharmaceutical company author (as the number of articles from companies at the tail end of our list is very small). While we have attempted to create a comparator with a globally representative group of research-intensive academic institutions, our design of a comparator set may not represent academic institutions in general because it includes only the largest academic institutions (in terms of publication counts) from each region.

We acknowledge the disparity in sample sizes between the pharmaceutical and academic datasets, which reflects the substantially higher publishing output of academic institutions. While this imbalance introduces potential for analytical bias and affects the stability and precision of percentages, it does not invalidate the descriptive analysis presented herein. Such biases are chiefly a concern for formal statistical analysis, which has not been undertaken here. Reducing the academic dataset would compromise representativeness, and random sampling within institutions is not currently supported on the Lens platform.

There is also overlap in our pharmaceutical company and academic institution analysis sets because most pharmaceutical company articles include academic authors (but not necessarily from the institutions in our academic institution analysis set). Hence, some individual articles may have been included in both analysis sets. Academic articles may also include pharmaceutical company authors (although in our analysis, over 90% did not). We also rely on accurate parsing of author affiliations by the Lens to identify pharmaceutical company and academic institution articles. Finally, there is a gray area consisting of articles that are based on studies initiated by academic institutions and funded by grants from pharmaceutical companies. Pharmaceutical companies, though providing financial support for such research, have no influence over publication decisions, such as target journals or whether to publish OA, and the articles typically do not have pharmaceutical company authors. Additionally, the funding provided usually does not cover publication costs such as APCs, and authors of these articles may face challenges to publishing OA.

### Future prospects

The OA publication rates for medical research articles are likely to continue to rise, both for pharmaceutical companies and academic institutions, but especially for pharmaceutical companies, which we believe will be beneficial to healthcare practitioners and patients. The continued increase in OA publication rates will likely be driven by growing awareness of the benefits of and options for OA, and by changes in policy at the journal and research funder (including pharmaceutical companies) levels. For example, journal policies that allow pharmaceutical company research to be published OA and under non-restrictive CC licenses could help incentivize authors of pharmaceutical company articles to choose OA publishing by giving them a wider range of target journals to choose from. The appetite among pharmaceutical company authors to publish their articles OA may be greater than the observed OA publication rates, given that several journals frequently targeted by pharmaceutical company research do not allow industry-funded research to be published OA (Lancet 2024; NEJM 2024) Research funders could introduce OA mandates that oblige authors to publish OA and use certain licenses.

## CONCLUSIONS

Our dashboard enables ongoing monitoring, and any individual is free to use this tool in the future. Measurement and competition are strong drivers for change and, although we have used pooled data here, it is possible to extract data for individual pharmaceutical companies and academic institutions. Combined with the ability to compare the OA publication rates by therapy area, the up-to-date tracking of OA publishing performance provided by the dashboard will help to identify and address concerns and bottlenecks that may hinder the ongoing adoption of OA publication practices. We welcome more research on this topic, and ever greater openness in the publication and dissemination of medical research.

## ACKNOWLEDGMENTS

The authors are grateful to the following people for their support in this work: Hollie Rawlings, MSc, of Oxford PharmaGenesis, Oxford, UK, for providing technical assistance with the dashboard; Aaron Ballagh, BASc, of the Lens, Canberra, Australia, for providing technical support with the Lens API and dashboard; Caitlin Edgell, PhD (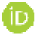: https://orcid.org/0000-0003-0448-122X), and Sophie Nobes, BSc (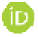: https://orcid.org/0009-0000-7816-0862), of Oxford PharmaGenesis, Oxford, UK, for providing medical writing support in accordance with Good Publication Practice 2022 (GPP 2022) guidelines (https://www.ismpp.org/gpp-2022); Velissaria Vanna, PhD, of Oxford PharmaGenesis, Oxford, UK, for providing editorial support; Kim Johnson, MScPH, of Oxford PharmaGenesis, Oxford, UK, for providing project management; and the Members, Supporters, Advisers and Followers of Open Pharma.

