## Supplementary material for "Benchmarking of open access publication rates for the pharmaceutical industry and research-intensive academic institutions: development and demonstration of a live, open monitoring tool"

### Plain Language Summary

Biotechnology and pharmaceutical companies develop and deliver most of the world’s new medicines. To make sure that important information about these medicines is shared transparently with people across the world, pharmaceutical companies are more and more often choosing to publish their research open access. Open access is a form of publishing that allows anyone to read and, in some cases, reuse the information in research articles for free. The proportion of articles published open access in medical journals is growing every year. However, we don’t know how the open access publication rates compare between medical articles published by pharmaceutical companies and medical articles published by academic institutions. Knowing this could help us understand how open access publication practices are changing over time and the impact of these changes.

We made a free online dashboard that tracks how many medical research articles from the top 40 pharmaceutical companies and 40 top universities are open access. The dashboard shows the number of medical research articles published across three time frames: one to two years before the dashboard is used; up to one year before the dashboard is used; and up to ten years before the dashboard is used. We downloaded data from the dashboard between 24 July and 4 August 2023.

We found that between 24 July 2021 and 24 July 2022 77% of pharmaceutical company research articles and 70% of academic institution research articles were published open access. Between 24 July 2013 and 24 July 2023, open access publication rates grew each year for both pharmaceutical company and academic institution research articles, but the rates grew faster for pharmaceutical companies.

Our results are similar to those from other studies of open access publication rates. We encourage people to explore our free online live report and share their results publicly. The live report can be accessed using the following link: <https://www.lens.org/lens/report/view/Open-access-dashboard/14572/page/14573>.

### Table S1 List of pharmaceutical companies included in this analysis (10-year timeframe).

| **Pharmaceutical company** | **HQ location** | **Total number of medical articles** | **Non-OA** | **OA** |
| --- | --- | --- | --- | --- |
| **AbbVie** | United States | 4077 | 1349 | 2728 |
| **Amgen** | United States | 4162 | 1644 | 2518 |
| **Astellas Pharma**† | Japan | 1595 | 685 | 910 |
| **AstraZeneca***† | United Kingdom | 8312 | 3052 | 5260 |
| **Bausch Health** | Canada | 128 | 38 | 90 |
| **Bayer** | Germany | 3946 | 1522 | 2424 |
| **Biogen** | United States | 1618 | 633 | 985 |
| **Boehringer Ingelheim***† | Germany | 3905 | 1381 | 2524 |
| **Bristol Myers Squibb***† | United States | 4726 | 1925 | 2801 |
| **CSL** | Australia | 626 | 172 | 454 |
| **CSPC Pharmaceutical Group** | Hong Kong | 11 | 5 | 6 |
| **Daiichi Sankyo** | Japan | 1457 | 615 | 842 |
| **Eisai** | Japan | 1366 | 595 | 771 |
| **Eli Lilly**† | United States | 5304 | 1801 | 3503 |
| **Fresenius** | Germany | 1216 | 373 | 843 |
| **Galápagos***† | Belgium | 214 | 39 | 175 |
| **Gilead Sciences***† | United States | 1928 | 691 | 1237 |
| **GlaxoSmithKline***† | United Kingdom | 7137 | 1939 | 5198 |
| **Ipsen***† | France | 638 | 250 | 388 |
| **Jazz Pharmaceuticals** | Ireland | 411 | 113 | 298 |
| **Jiangsu Hengrui Medicine** | China | 140 | 86 | 54 |
| **Johnson & Johnson***† | United States | 8259 | 2977 | 5282 |
| **Kyowa Kirin** | Japan | 353 | 97 | 256 |
| **Merck KGaA** | Germany | 8376 | 3492 | 4884 |
| **Moderna** | United States | 222 | 48 | 174 |
| **Merck & Co.** | United States | 1410 | 419 | 991 |
| **Novartis***† | Switzerland | 9781 | 4075 | 5706 |
| **Novo Nordisk***† | Denmark | 3259 | 743 | 2516 |
| **Otsuka** | Japan | 855 | 266 | 589 |
| **Pfizer***† | United States | 9014 | 3169 | 5845 |
| **Regeneron** | United States | 1500 | 467 | 1033 |
| **Roche***† | Switzerland | 10 229 | 4196 | 6033 |
| **Sanofi** | France | 4650 | 1238 | 3412 |
| **Sino Biopharmaceutical** | China | 0 | 0 | 0 |
| **Sumitomo Pharma** | Japan | 203 | 69 | 134 |
| **Takeda***† | Japan | 3819 | 1157 | 2662 |
| **Teva Pharmaceuticals** | Israel | 401 | 210 | 191 |
| **UCB***† | Belgium | 1485 | 491 | 994 |
| **Vertex Pharmaceuticals** | United States | 338 | 125 | 213 |
| **Viatris** | United States | 80 | 12 | 68 |

**Notes.**

*Open Pharma Member or Supporter company in Q3 2023. †Open Pharma Member or Supporter company in Q4 2024.

Abbreviations: HQ, headquarters; OA, open access.

### Table S2 List of academic institutions included in this analysis (10-year timeframe).

| **Academic institution** | **Region** | **Total number of medical articles** | **Non-OA** | **OA** |
| --- | --- | --- | --- | --- |
| **Capital Medical University** | Asia | 40 420 | 17 625 | 22 795 |
| **Duke University** | North America | 45 847 | 17 354 | 28 493 |
| **Erasmus University Rotterdam** | Europe | 23 032 | 6881 | 16 151 |
| **Freie Universität Berlin** | Europe | 7680 | 2057 | 5623 |
| **Fudan University** | Asia | 38 663 | 15 817 | 22 846 |
| **Harvard University** | North America | 148 452 | 55 078 | 93 374 |
| **Heidelberg University** | Europe | 22 089 | 8379 | 13710 |
| **Huazhong University of Science and Technology** | Asia | 25 298 | 9439 | 15 859 |
| **Humboldt-Universität zu Berlin** | Europe | 7396 | 1908 | 5488 |
| **Imperial College London** | Europe | 38 992 | 10 248 | 28 744 |
| **Johns Hopkins University** | North America | 76 044 | 26 739 | 49 305 |
| **Karolinska Institutet** | Europe | 44 992 | 12 479 | 32 513 |
| **King's College London** | Europe | 36 634 | 9267 | 27 367 |
| **Nanjing Medical University** | Asia | 26 970 | 10 789 | 16 181 |
| **Peking Union Medical College** | Asia | 46 241 | 19 252 | 26 989 |
| **Peking University** | Asia | 40 010 | 17 006 | 23 004 |
| **Seoul National University** | Asia | 39 041 | 14 630 | 24 411 |
| **Shanghai Jiao Tong University** | Asia | 43 704 | 18 052 | 25 652 |
| **Sichuan University** | Asia | 31 250 | 13 170 | 18 080 |
| **Stanford University** | North America | 52 168 | 20 120 | 32 048 |
| **Sun Yat-sen University** | Asia | 37 868 | 14 775 | 23 093 |
| **Tel Aviv University** | Middle East | 21 816 | 11 002 | 10 814 |
| **University of Paris** | Europe | 35 751 | 13 508 | 22 243 |
| **University College London** | Europe | 59 877 | 13 703 | 46 174 |
| **University of Amsterdam** | Europe | 26 344 | 7100 | 19 244 |
| **University of California, San Francisco** | North America | 59 494 | 20 184 | 39 310 |
| **University of Cambridge** | Europe | 32 680 | 8241 | 24 439 |
| **University of Cape Town** | Africa | 14 481 | 3200 | 11 281 |
| **University of Copenhagen** | Europe | 32 626 | 11 703 | 20 923 |
| **University of Melbourne** | Oceania | 42 212 | 14 903 | 27 309 |
| **University of Michigan** | North America | 55 264 | 19 829 | 35 435 |
| **University of Milan** | Europe | 34 921 | 13 121 | 21 800 |
| **University of Oxford** | Europe | 42 041 | 8559 | 33 482 |
| **University of Pennsylvania** | North America | 59 691 | 23 911 | 35 780 |
| **University of São Paulo** | South America | 55 477 | 22 113 | 33 364 |
| **University of Sydney** | Oceania | 47 360 | 19 156 | 28 204 |
| **University of Toronto** | North America | 72 473 | 29 790 | 42 683 |
| **University of Zurich** | Europe | 25 499 | 7450 | 18 049 |
| **Utrecht University** | Europe | 25 210 | 6803 | 18 407 |
| **Zhejiang University** | Asia | 33 465 | 12 765 | 20 700 |

Abbreviation: OA, open access.

### Table S3 Regional composition of academic institutions selected for this analysis (data from the CWTS Leiden Ranking 2022).

|  | **All academic institutions** | | | **Academic institutions selected for analysis** | | |
| --- | --- | --- | --- | --- | --- | --- |
| **Region** | **Total academic institutions (n)** | **Total articles** | **% of total** | **Selected academic institutions (n)** | **Total articles** | **% of total** |
| **Africa** | 16 | 26 686 | 0.6 | 1 | 5080 | 0.7 |
| **Asia** | 495 | 1 023 041 | 24.8 | 11 | 184 920 | 23.9 |
| **Europe** | 441 | 1 434 347 | 34.8 | 16 | 266 866 | 34.4 |
| **Middle East** | 39 | 72 199 | 1.8 | 1 | 11 659 | 1.5 |
| **North America** | 240 | 1 294 201 | 31.4 | 8 | 248 738 | 32.1 |
| **Oceania** | 41 | 177 318 | 4.3 | 2 | 38 731 | 5.0 |
| **South America** | 46 | 96 154 | 2.3 | 1 | 18 863 | 2.4 |
| **Grand Total** | 1318 | 4 123 944 | 100.0 | 40 | 774 857 | 100.0 |

Abbreviation: CWTS, The Centre for Science and Technology Studies.

### Table S4 Full list of ‘field of study’ tags for the top 10 therapy areas selected for this analysis.

| **Therapy area** | **Field of study tags** |
| --- | --- |
| **Cardiovascular** | "Cardiology" OR "Heart failure" OR "Atrial fibrillation" OR "Myocardial infarction" OR "Stroke" OR "Blood pressure" OR "Acute coronary syndrome" OR "Coronary artery disease" OR "Thrombosis" OR "QT interval" OR "Heart rate" OR "Heart failure with preserved ejection fraction" OR "Cardiomyopathy" |
| **Endocrine and metabolic disorders** | "Endocrinology" OR "Diabetes mellitus" OR "Type 2 diabetes" OR "Insulin" OR "Type 2 diabetes mellitus" OR "Hypoglycemia" OR "Type 1 diabetes" OR "Insulin resistance" OR "Dyslipidemia" OR "Diabetic retinopathy" OR "Diabetic nephropathy" OR "Endocrine system" |
| **Gastroenterology** | "Gastroenterology" OR "Ulcerative colitis" OR "Inflammatory bowel disease" OR "Crohn's disease" |
| **Genitourinary disorders (including nephrology)** | "Urology" OR "Renal function" OR "Kidney disease" OR "Kidney" OR "Urinary system" OR "Urine" OR "End stage renal disease" OR "Acute kidney injury" |
| **Immunology and inflammation** | "Immunology" OR "Rheumatoid arthritis" OR "Immune system" OR "Inflammation" OR "Psoriatic arthritis" OR "Psoriasis" OR "Rheumatology" OR "Arthritis" OR "T cell" OR "Ankylosing spondylitis" OR "Cytotoxic T cell" OR "Proinflammatory cytokine" OR "Autoimmune disease" |
| **Microbiology (including infections)** | "Virology" OR "Virus" OR "Microbiology" OR "Infectious disease (medical specialty)" OR "Viral load" OR "Pneumonia" OR "Human immunodeficiency virus (HIV)" OR "Hepatitis C virus" OR "Hepatitis C" OR "Immunization" OR "Bacteria" OR "Tuberculosis" OR "Antibiotic resistance" OR "Hepatitis B virus" |
| **Oncology** | "Oncology" OR "Cancer" OR "Cancer research" OR "Chemotherapy" OR "Breast cancer" OR "Lung cancer" OR "Prostate cancer" OR "Pembrolizumab" OR "Multiple myeloma" OR "Colorectal cancer" OR "Neutropenia" OR "Metastatic breast cancer" OR "Lymphoma" OR "Myeloid leukemia" OR "Melanoma" OR "Renal cell carcinoma" OR "Leukemia" OR "Chronic lymphocytic leukemia" OR "Hepatocellular carcinoma" OR "Febrile neutropenia" OR "Carcinoma" OR "Ovarian cancer" OR "Response Evaluation Criteria in Solid Tumors" OR "Adenocarcinoma" OR "Follicular lymphoma" OR "Diffuse large B-cell lymphoma" OR "Cancer cell" OR "non-small cell lung cancer (NSCLC)" OR "Triple-negative breast cancer" OR "Pancreatic cancer" OR "Cancer immunotherapy" OR "Bladder cancer" OR "Cervical cancer" OR "Head and neck squamous-cell carcinoma" OR "Metastatic Urothelial Carcinoma" OR "Surgical oncology" OR "Head and neck cancer" OR "Carcinogenesis" |
| **Pharmacology** | "Pharmacology" OR "Pharmacokinetics" OR "Chemistry" OR "Biochemistry" OR "Pharmacodynamics" OR "In vitro" OR "Pharmacotherapy" OR "Clinical pharmacology" |
| **Psychiatry** | "Psychiatry" OR "Psychology" OR "Psychological intervention" OR "Dementia" OR "Clinical psychology" OR "Anxiety" OR "Major depressive disorder" OR "Schizophrenia" OR "Alzheimer's disease" |
| **Respiratory** | "COPD" OR "Asthma" OR "Lung" OR "Idiopathic pulmonary fibrosis" OR "Respiratory system" OR "Spirometry" OR "Pulmonary disease" OR "Respiratory tract infections" |

### Table S5 OA license categories used in this analysis.

| **OA license category** | **License types included*** |
| --- | --- |
| **Unrestricted OA** | CC0, Public Domain, CC BY |
| **Restricted OA** | CC BY-NC, CC BY-NC-ND, CC BY-NC-SA, CC BY-SA |
| **Publisher-specific** | Publisher-specific, Elsevier, ACS |
| **Unknown** | Implied-OA, Unknown |

**Notes.**

*Details of each license type can be found on the website of the Lens.^38^

Abbreviations: ACS, American Chemical Society; CC0, Creative Commons Zero; CC BY, Creative Commons Attribution; CC BY-NC, Creative Commons Attribution-NonCommercial; CC BY-NC-SA, Creative Commons Attribution-NonCommercial-ShareAlike; CC BY-ND, Creative Commons Attribution-NoDerivs; CC BY-SA, Creative Commons Attribution-ShareAlike; OA, open access.

### Table S6 Sen’s slope calculation of OA trends for pharmaceutical company and academic institution medical articles over 10 years (2013–2023).

|  | **Pharmaceutical company** | **Academic institution** |
| --- | --- | --- |
| **Change in overall percentage (%) OA** | | |
| Overall | 3.1 | 1.6 |
| **Change in percentage OA (%) across therapy areas** | | |
| Cardiovascular | 2.8 | 1.5 |
| Endocrinology | 2.7 | 1.0 |
| Gastroenterology | 4.1 | 2.0 |
| Genitourinary disorders | 3.5 | 1.3 |
| Immunology | 3.6 | 1.9 |
| Microbiology | 3.2 | 1.7 |
| Oncology | 2.8 | 1.5 |
| Pharmacology | 3.9 | 1.6 |
| Psychiatry | 3.4 | 1.3 |
| Respiratory | 3.9 | 2.3 |
| **Change in overall percentage (%) OA model*** | | |
| Bronze | –0.6 | –0.9 |
| Gold | 1.4 | 2.3 |
| Green | –2.2 | –2.6 |
| Hybrid | 1.1 | 0.9 |
| Unknown | 0.0 | 0.2 |

**Notes.**

*Percentage OA model was calculated based on total number of OA articles, not total article count.

Abbreviation: OA, open access.

### Table S7 Number of pharmaceutical company and academic institution medical articles for selected therapy areas (primary analysis, articles 12–24 months old).

| **Therapy areas** | **Pharmaceutical company** | **Academic institution** |
| --- | --- | --- |
| **All medical articles, n (%)** | 13 107 (100.0) | 184 672 (100.0) |
| **Cardiovascular, n (%)** | 918 (7.0) | 15 824 (8.6) |
| **Endocrinology, n (%)** | 1324 (10.1) | 15 892 (8.6) |
| **Gastroenterology, n (%)** | 1496 (11.4) | 10 276 (5.6) |
| **Genitourinary disorders (including nephrology), n (%)** | 668 (5.1) | 6249 (3.4) |
| **Immunology, n (%)** | 2468 (18.8) | 22 471 (12.2) |
| **Microbiology (including infections), n (%)** | 1323 (10.1) | 19 641 (10.6) |
| **Oncology, n (%)** | 4012 (30.6) | 35 223 (19.1) |
| **Pharmacology, n (%)** | 3109 (23.7) | 27 743 (15.0) |
| **Psychiatry, n (%)** | 1399 (10.7) | 38 107 (20.6) |
| **Respiratory, n (%)** | 642 (4.9) | 5055 (2.7) |

### Table S8 Journals with the highest number of pharmaceutical company and academic institution medical articles in each therapy area (primary analysis, articles 12–24 months old). Shaded rows indicate journals that appear in the top 5 in both groups.

| **Top pharmaceutical company journals** | **Number of articles** | **% of total articles** | **% of journal articles that are OA** | **Top academic institution journals** | **Number of articles** | **% of total articles** | **% of journal articles that are OA** |
| --- | --- | --- | --- | --- | --- | --- | --- |
| **Cardiovascular** | | | | | | | |
| ***European Heart Journal*** | 43 | 0.3 | 74.4 | ***Frontiers in Cardiovascular Medicine*** | 724 | 0.4 | 100.0 |
| ***European Journal of Heart Failure*** | 23 | 0.2 | 87.0 | ***Journal of the American Heart Association*** | 278 | 0.2 | 100.0 |
| ***Circulation*** | 21 | 0.2 | 81.0 | ***SSRN Electronic Journal*** | 259 | 0.1 | 7.7 |
| ***Journal of the American Heart Association*** | 20 | 0.2 | 100.0 | ***Frontiers in Neurology*** | 205 | 0.1 | 100.0 |
| ***ESC Heart Failure*** | 17 | 0.1 | 100.0 | ***Scientific Reports*** | 205 | 0.1 | 100.0 |
| **Endocrinology** | | | | | | | |
| ***Diabetes, Obesity & Metabolism*** | 83 | 0.6 | 83.1 | ***Frontiers in Endocrinology*** | 439 | 0.2 | 100.0 |
| ***Diabetes Therapy: Research, Treatment and Education of Diabetes and Related Disorders*** | 52 | 0.4 | 100.0 | ***The Journal of Clinical Endocrinology and Metabolism*** | 299 | 0.2 | 63.2 |
| ***Advances In Therapy*** | 19 | 0.1 | 78.9 | ***Scientific Reports*** | 231 | 0.1 | 100.0 |
| ***International Journal of Molecular Sciences*** | 18 | 0.1 | 100.0 | ***SSRN Electronic Journal*** | 227 | 0.1 | 10.1 |
| ***Nephrology Dialysis Transplantation*** | 18 | 0.1 | 27.8 | ***Nutrients*** | 215 | 0.1 | 100.0 |
| **Gastroenterology** | | | | | | | |
| ***Journal of Clinical Oncology*** | 185 | 1.4 | 0.5 | ***Journal of Clinical Oncology*** | 464 | 0.3 | 1.3 |
| ***Journal of Crohn's and Colitis*** | 93 | 0.7 | 100.0 | ***Frontiers in Oncology*** | 220 | 0.1 | 100.0 |
| ***HemaSphere*** | 70 | 0.5 | 100.0 | ***Frontiers in Immunology*** | 163 | 0.1 | 100.0 |
| ***Clinical Cancer Research*** | 41 | 0.3 | 80.5 | ***Frontiers in Medicine*** | 155 | 0.1 | 100.0 |
| ***Blood*** | 35 | 0.3 | 97.1 | ***Journal of Crohn's and Colitis*** | 146 | 0.1 | 100.0 |
| **Genitourinary disorders (including nephrology)** | | | | | | | |
| ***Journal of Clinical Oncology*** | 68 | 0.5 | 0.0 | ***Food and Chemical Toxicology*** | 172 | 0.1 | 0.0 |
| ***Nephrology Dialysis Transplantation*** | 42 | 0.3 | 33.3 | ***Journal of Clinical Oncology*** | 160 | 0.1 | 0.6 |
| ***Kidney International Reports*** | 17 | 0.1 | 100.0 | ***Frontiers in Medicine*** | 118 | 0.1 | 100.0 |
| ***Diabetes, Obesity & Metabolism*** | 16 | 0.1 | 87.5 | ***Nephrology Dialysis Transplantation*** | 93 | 0.1 | 30.1 |
| ***Nephrology, Dialysis, Transplantation*** | 11 | 0.1 | 90.9 | ***Scientific Reports*** | 85 | 0.0 | 100.0 |
| **Immunology** | | | | | | | |
| ***Annals of the Rheumatic Diseases*** | 145 | 1.1 | 98.6 | ***Frontiers in Immunology*** | 1108 | 0.6 | 100.0 |
| ***Journal of Clinical Oncology*** | 51 | 0.4 | 0.0 | ***SSRN Electronic Journal*** | 419 | 0.2 | 11.0 |
| ***Vaccine*** | 47 | 0.4 | 70.2 | ***Frontiers in Oncology*** | 246 | 0.1 | 100.0 |
| ***Advances in Therapy*** | 44 | 0.3 | 95.5 | ***Scientific Reports*** | 234 | 0.1 | 100.0 |
| ***Open Forum Infectious Diseases*** | 44 | 0.3 | 100.0 | ***Annals of the Rheumatic Diseases*** | 231 | 0.1 | 98.7 |
| **Microbiology (including infections)** | | | | | | | |
| ***Open Forum Infectious Diseases*** | 132 | 1.0 | 100.0 | ***SSRN Electronic Journal*** | 761 | 0.4 | 16.8 |
| ***Human Vaccines & Immunotherapeutics*** | 40 | 0.3 | 100.0 | ***PloS One*** | 311 | 0.2 | 100.0 |
| ***Vaccine*** | 37 | 0.3 | 67.6 | ***Frontiers in Immunology*** | 255 | 0.1 | 100.0 |
| ***Infectious Diseases and Therapy*** | 32 | 0.2 | 100.0 | ***Scientific Reports*** | 240 | 0.1 | 100.0 |
| ***SSRN Electronic Journal*** | 28 | 0.2 | 21.4 | ***Clinical Infectious Diseases*** | 231 | 0.1 | 85.3 |
| **Oncology** | | | | | | | |
| ***Journal of Clinical Oncology*** | 717 | 5.5 | 0.3 | ***Journal of Clinical Oncology*** | 1851 | 1.0 | 1.1 |
| ***Blood*** | 329 | 2.5 | 100.0 | ***Frontiers in Oncology*** | 1529 | 0.8 | 100.0 |
| ***HemaSphere*** | 184 | 1.4 | 100.0 | ***Blood*** | 940 | 0.5 | 97.8 |
| ***Cancer Research*** | 157 | 1.2 | 0.0 | ***Cancers*** | 782 | 0.4 | 100.0 |
| ***Clinical Cancer Research*** | 91 | 0.7 | 83.5 | ***Cancer Research*** | 573 | 0.3 | 0.0 |
| **Pharmacology** | | | | | | | |
| ***Journal of Clinical Oncology*** | 102 | 0.8 | 0.0 | ***SSRN Electronic Journal*** | 781 | 0.4 | 7.4 |
| ***Journal of Medicinal Chemistry*** | 62 | 0.5 | 24.2 | ***Frontiers in Pharmacology*** | 535 | 0.3 | 100.0 |
| ***Clinical and Translational Science*** | 55 | 0.4 | 100.0 | ***Frontiers in Immunology*** | 382 | 0.2 | 100.0 |
| ***Clinical Pharmacology and Therapeutics*** | 51 | 0.4 | 58.8 | ***Scientific Reports*** | 334 | 0.2 | 100.0 |
| ***HemaSphere*** | 41 | 0.3 | 100.0 | ***Journal of Clinical Oncology*** | 297 | 0.2 | 0.7 |
| **Psychiatry** | | | | | | | |
| ***Alzheimer's & Dementia*** | 59 | 0.5 | 98.3 | ***SSRN Electronic Journal*** | 776 | 0.4 | 11.9 |
| ***Sleep*** | 22 | 0.2 | 68.2 | ***BMJ Open*** | 687 | 0.4 | 100.0 |
| ***Neurology and Therapy*** | 21 | 0.2 | 100.0 | ***International Journal of Environmental Research and Public Health*** | 676 | 0.4 | 100.0 |
| ***Advances in Therapy*** | 19 | 0.1 | 89.5 | ***Frontiers in Psychiatry*** | 598 | 0.3 | 100.0 |
| ***BMC Psychiatry*** | 19 | 0.1 | 100.0 | ***Alzheimer's & Dementia*** | 534 | 0.3 | 99.6 |
| **Respiratory** | | | | | | | |
| ***International Journal of Chronic Obstructive Pulmonary Disease*** | 30 | 0.2 | 100.0 | ***SSRN Electronic Journal*** | 112 | 0.1 | 9.8 |
| ***The Journal of Allergy and Clinical Immunology. In Practice*** | 27 | 0.2 | 77.8 | ***Frontiers in Immunology*** | 94 | 0.1 | 100.0 |
| ***Respiratory Research*** | 24 | 0.2 | 100.0 | ***American Journal of Respiratory and Critical Care Medicine*** | 80 | 0.0 | 75.0 |
| ***ERJ Open Research*** | 18 | 0.1 | 100.0 | ***The European Respiratory Journal*** | 79 | 0.0 | 59.5 |
| ***The European Respiratory Journal*** | 18 | 0.1 | 72.2 | ***Scientific Reports*** | 76 | 0.0 | 100.0 |

Abbreviation: OA, open access.

### Table S9 Sensitivity analysis of OA model (color) used in pharmaceutical company, all academic institution and North American and European academic institution medical articles.

| **OA model (color)*** | **Pharmaceutical companies** | **All academic institutions** | **North American  and European  academic institutions** |
| --- | --- | --- | --- |
| **Primary analysis (articles 12–24 months old†)** | | | |
| Bronze, n (%) | 1 792 (12.4) | 16 336 (8.4) | 12 885 (10.7) |
| Gold, n (%) | 5 229 (36.3) | 80 342 (41.4) | 42 429 (35.1) |
| Hybrid, n (%) | 3 122 (21.7) | 21 715 (11.2) | 17 478 (14.5) |
| Green, n (%) | 835 (5.8) | 16 906 (8.7) | 13 638 (11.3) |
| Non-OA, n (%) | 3 251 (22.6) | 53 651 (27.6) | 30 283 (25.1) |
| Unknown, n (%) | 171 (1.2) | 5 108 (2.6) | 4 066 (3.4) |

**Notes.**

*Percentages indicate percentage of total counts for the time frame and comparison group. †Article count for the 12-month period 12–24 months prior to original data extraction (24 July 2021–24 July 2022). This data set was downloaded on 12 December 2024.

Abbreviation: OA, open access.

### Table S10 Sensitivity analysis of overall OA/non-OA status and license categories used in pharmaceutical company, all academic institution and North American and European academic institution medical articles.

| **OA status and licence category*** | **Pharmaceutical companies** | **All academic institutions** | **North American  and European  academic institutions** |
| --- | --- | --- | --- |
| **Primary analysis (articles 12–24 months old†)** | | | |
| Total, n (%) | 14 450 (100) | 195 457 (100) | 121 396 (100) |
| Non-OA, n (%) | 3290 (22.8) | 55 196 (28.2) | 30 757 (25.3) |
| OA (any licence), n (%) | 11 160 (77.2) | 140 261 (71.8) | 90 639 (74.7) |
| OA, unrestricted licence, n (%) | 3210 (22.2) | 59 698 (30.5) | 33 497(27.6) |
| OA, restricted licence, n (%) | 4963 (34.3) | 38 627 (19.8) | 24 667 (20.3) |
| OA, publisher licence, n (%) | 113 (0.8) | 2615 (1.3) | 2187 (1.8) |
| OA, unknown licence, n (%) | 2870 (19.9) | 39 138 (20.0) | 30 187 (24.9) |

**Notes.**

*Percentages indicate percentage of total counts for the time frame and comparison group. †Article count for the 12-month period 12–24 months prior to original data extraction (24 July 2021–24 July 2022). This data set was downloaded on 20 November 2024.

Abbreviation: OA, open access.
